# Human V4 size predicts crowding distance

**DOI:** 10.1101/2024.04.03.587977

**Authors:** Jan W. Kurzawski, Brenda S. Qiu, Najib J. Majaj, Noah Benson, Denis G. Pelli, Jonathan Winawer

## Abstract

Visual recognition is limited by both object size (acuity) and spacing. The spacing limit, called “crowding”, is the failure to recognize an object in the presence of other objects. Here, we take advantage of individual differences in crowding to investigate its biological basis. Crowding distance, the minimum object spacing needed for recognition, varies 2-fold among healthy adults. We test the conjecture that this variation in psychophysical crowding distance is due to variation in cortical map size. To test this, we made paired measurements of brain and behavior in 50 observers. We used psychophysics to measure crowding distance and calculate *λ*, the number of letters that fit into each observer’s visual field without crowding. In the same observers, we used fMRI to measure the surface area *A* (mm²) of retinotopic maps V1, V2, V3, and V4. Across observers, *λ* is proportional to the surface area of V4 but is uncorrelated with the surface area of V1 to V3. The proportional relationship of *λ* to area of V4 indicates conservation of *cortical crowding distance* across individuals: letters can be recognized if they are spaced by at least 1.4 mm on the V4 map, irrespective of map size and psychophysical crowding distance. We conclude that the size of V4 predicts the spacing limit of visual perception.

## Introduction

Visual recognition is limited by two independent scale factors: the size and spacing of objects^1^. The size limit is understood but the spacing limit is not. The size limit (acuity) is set by properties of the eye, including optical quality and the sampling density of the retinal cell mosaics^2^. However, even when object size is well above the acuity limit, recognition fails if objects are insufficiently spaced (crowding; **Fig. 1**).

**Figure 1.**
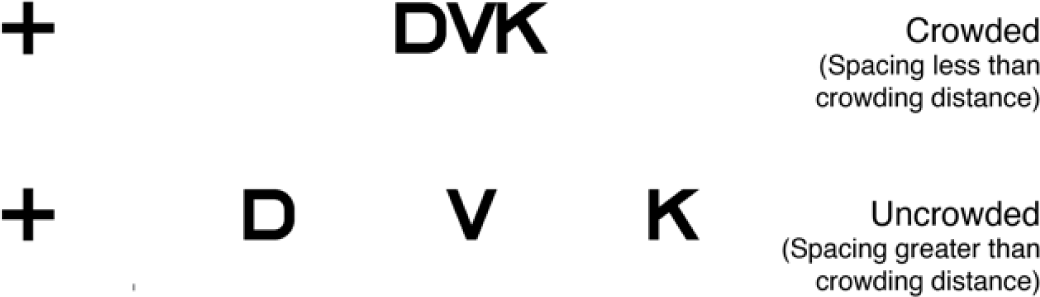
Example of crowding. Top: While fixating the cross, the letter V is unrecognizable (“crowded”) because the center-to-center letter spacing is less than the crowding distance. Bottom: The letter V is recognizable when the letter spacing is greater than the crowding distance.

Crowding limits performance of important everyday activities, including reading and search^3,4^. The severity of this limitation varies among healthy individuals and between groups: Crowding thresholds are elevated in several disorders, including dyslexia^5^, dyscalculia^6^, strabismic amblyopia^1^ and apperceptive agnosia^7^. Although studied for over a century^8^, the biological basis of crowding is largely unknown. Physiological studies have yielded conflicting claims about the anatomical site of crowding (**Table 1**). Perhaps the only certain conclusion is that crowding has a cortical basis, because it persists when target and clutter are presented separately to the two eyes^9,10^.

**Table 1.**
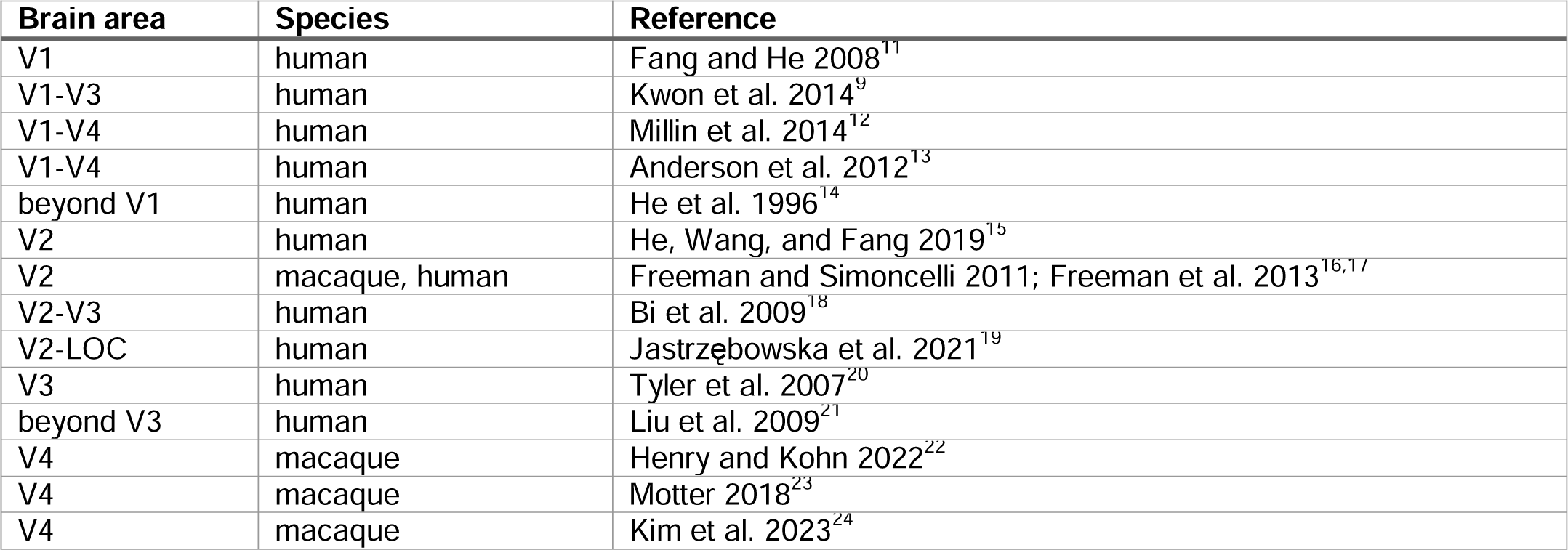
Physiology-based links between crowding and visual brain areas.

Here, we take an anatomical rather than physiological approach to study the basis of crowding. We do so by taking advantage of variation in crowding distance and cortical map size across observers.

Individual differences in crowding^25–28^ and map size^29^ are large even among young healthy observers. We conjecture that crowding results from limited neural resources at some processing stage, resembling the way acuity is limited by retinal cell density. Retinotopic maps in the human cortex are obvious candidates to mediate crowding. We ask whether the surface area of one or more retinotopic maps is associated with crowding distance variability across observers. Further, we test the more specific hypothesis that the *cortical crowding distance* (threshold object spacing on the cortical surface) is a constant for one or more retinotopic maps, despite large individual differences in psychophysical crowding distance and map size.

Some quantities are known to be conserved in cortical architecture (for review see ref [^30^]). For example, in visual cortex, V1 cortical magnification and population receptive field size both vary substantially across eccentricities and individuals, yet the product of these two measures, “the population point image”, is approximately constant^31^. This indicates that a point in the visual field is represented by the same amount of V1 cortex, irrespective of the observer and eccentricity. A second example is found in the mapping between visual areas: Receptive fields in macaque V4 grow with eccentricity when measured in degrees of visual field; however, within an individual animal, they maintain a constant size, independent of eccentricity, when projected onto the map of V1, showing constancy in intracortical sampling^32^. This implies that the way higher level brain areas sample V1 is relatively constant within an individual. The quantities that are conserved are likely to be important for explaining the functional organization of the nervous system.

Here we use the concept of cortical conservation to test the possibility that variation in psychophysical crowding distance is predicted by the surface area of retinotopic maps.

## Results

### Cortical conservation hypothesis

In each of 50 observers, we psychophysically measure the crowding distance with a letter identification task, and use fMRI to measure the surface area of retinotopic maps V1 to V4. Inspired by the letter acuity charts introduced by Anstis in 1974^33^, we calculate *λ*, the maximum number of letters that fit into an observer’s visual field without crowding (**Figure 2a**). The smaller the crowding distance the more letters fit in their visual field. Conservation of cortical crowding distance implies that variation in *λ* is entirely due to variation in surface area *A* of one or more maps, such that observers with larger maps can recognize proportionately more letters in their visual field (**Figure 2b**). We express this relationship as,

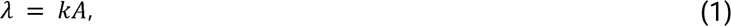

where *λ* (letters) and *A* (mm^2^) are vectors with one element per observer, and *k* is a constant scalar (letters per mm^2^ of cortex). Writing *k* as a constant expresses the hypothesis that variation in *λ* across observers is due only to variation in *A*.

**Figure 2.**
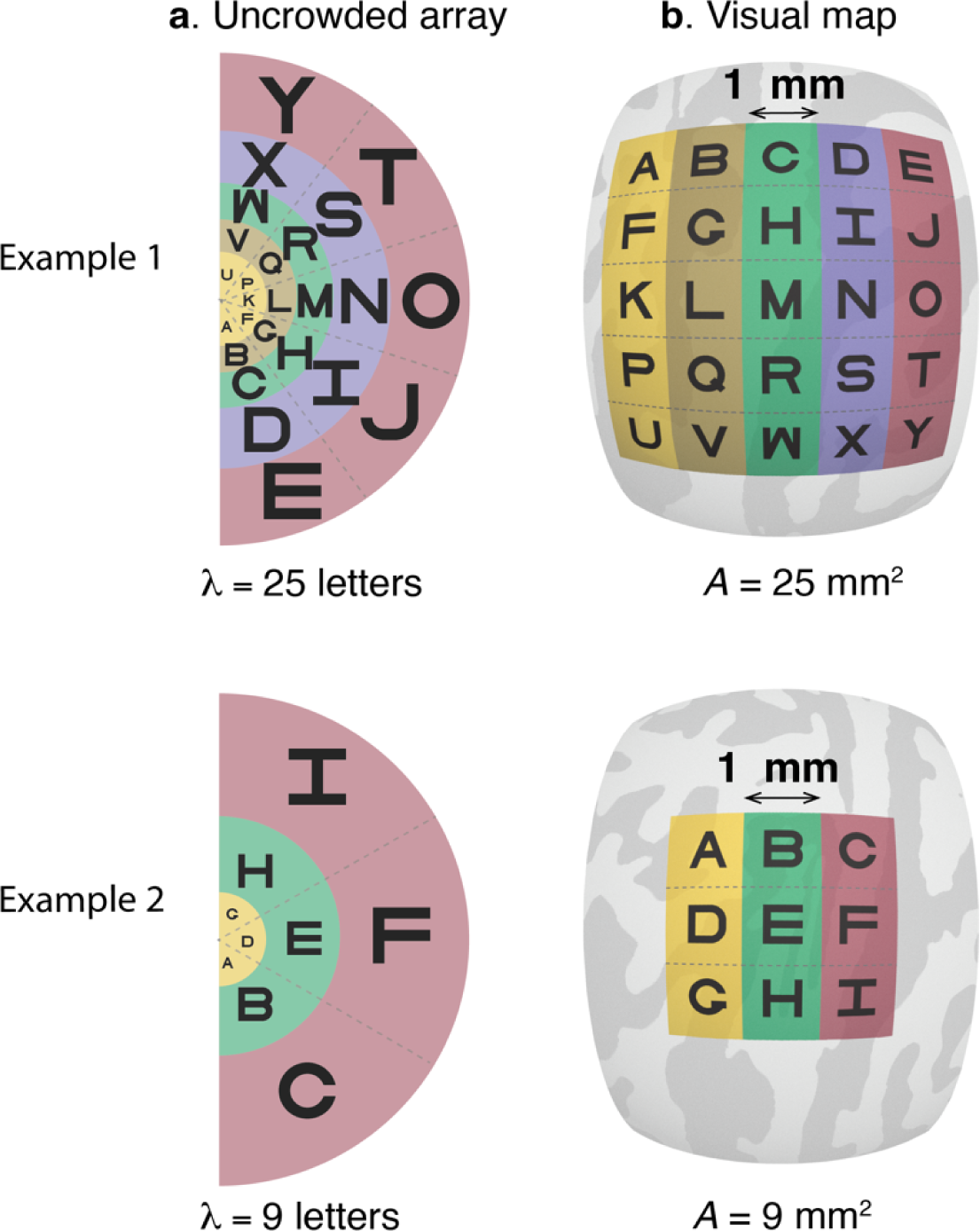
Conservation of cortical crowding distance. (a) Two hypothetical examples of uncrowded visual fields. In each case, the letters are spaced at the minimum distance necessary for successful recognition. In both examples, the spacing increases with eccentricity. The denser array in Example 1 indicates that this hypothetical observer is more tolerant to clutter (smaller crowding distance), and hence can fit more letters into the uncrowded visual field than observer 2 (*λ* = 25 vs *λ* = 9). (b) The projection of the letter arrays onto hypothetical retinotopic maps on the cortical surface. The greater *λ* for the first observer is explained by a larger cortical map, but with equal center-to-center letter spacing on the cortex. Hence the conservation conjecture predicts that the number of letters in the uncrowded field is proportional to surface area. The projections assume cortical conservation of crowding with a constant *k* = 1 letter / mm^2^.

Prior work observed that radial crowding distance, in degrees of visual angle, increases linearly with eccentricity (the Bouma law), and that the radial cortical magnification, measured in mm/deg, decreases inversely with eccentricity, such that the product of the two functions is roughly constant^34^ (but see ^35^). This amounts to conservation across eccentricity of the threshold center-to-center letter spacing on the cortical surface. Critically, this conservation applies equally to any retinotopic map where cortical magnification decreases inversely with eccentricity, without favoring any map for association with psychophysical crowding^34^. In contrast, conservation across individuals might favor a single map, enabling us to link crowding to a brain locus. Indeed, across individuals, the sizes of retinotopic maps vary largely independently, meaning that the size of one map (e.g., V1) only weakly predicts the size of others (e.g., V4), with the exception of adjacent maps such as V1 and V2^29,36^. Here we test each retinotopic map for conservation of crowding distance across individuals, with reason to expect that it cannot be conserved in multiple maps.

### Individual differences in crowding and map size

Consider two actual observers, one with a large crowding distance and one with a small crowding distance (**Figure 3a**). As expected, for both observers crowding distance grows with eccentricity. Dividing the crowding distance by eccentricity gives the Bouma factor, which is 0.22±0.03 for observer 1 and 0.34±0.05 for observer 2 [mean±SD_test-retest_]. Using the Bouma factors, we then compute, the number of letters that can fit into the visual field spaced at their psychophysically measured crowding distances (**Figure 3b**). This value is much larger for Observer 1 than 2 (748 vs. 302). We next estimate map surface area for each observer using retinotopy (**Figure 3c**). Observer 1’s V4 surface area is twice Observer 2’s, whereas V1 to V3 surface areas are about the same for the two observers (**Table 2**).

**Figure 3.**
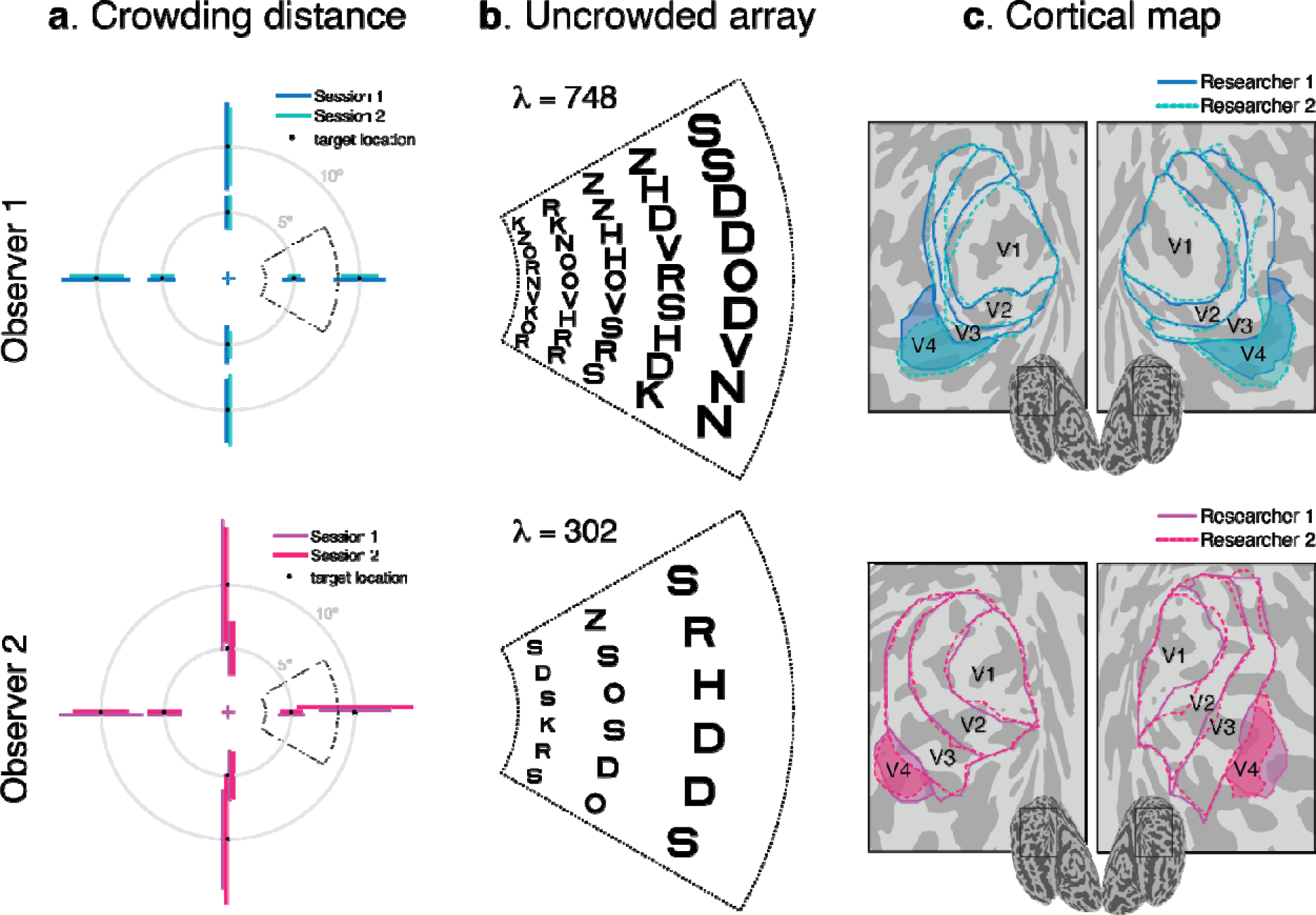
Individual differences in crowding and map size. (a) Crowding distances for Observers 1 and 2. The crowding distance was measured radially at two eccentricities, 5° and 10°, in the four cardinal directions, twice at each location. The eight target locations are indicated by black dots. Crowding thresholds are displayed as horizontal and vertical lines. The endpoints of the lines indicate the neighboring letter locations at threshold spacing. Observer 2’s lines are slightly offset to avoid overlap. Dotted sectors represent the part of the visual field shown in Panel b. (b) For each observer, we show a portion of the visual field with letters spaced according to their Bouma factor. The portion of the visual field shown is 3° to 8.5° eccentricity and ±25° polar angle. The values count letters for the whole visual field up to 10° eccentricity, the extent of our psychophysics and retinotopy. (c) Retinotopic map boundaries (V1 to V4) from both hemispheres of the same two observers. Visual maps were independently drawn by two researchers (dashed and solid lines). Polar angle and eccentricity maps for the two observers are shown in **Supplementary Figure 1**. Maps for all 49 participants are included as Supplementary Data 2.

**Table 2.** Number of letters in uncrowded visual field and surface area *A* in two observers. The numbers in the table indicate each observer’s and each observer’s surface area A, in mm^2^ of the bilateral retinotopic maps and the ratios between observers. Surface areas are averaged across the two researchers (**Figure 3C**). The two observers are the same as the two in **Figure 3**. Data for all observers are provided in Supplementary Data 1.

|  |  | A (mm<sup>2</sup>) |  |  |  |
| --- | --- | --- | --- | --- | --- |
|  |  | V1 | V2 | V3 | V4 |
| Observer 1 | 748 | 3052 | 2767 | 2053 | 1483 |
| Observer 2 | 302 | 2580 | 2815 | 1907 | 618 |
| Ratio | 2.5 | 1.2 | 1.0 | 1.1 | 2.4 |

### Test-retest reliability of and *A*

Testing the conservation conjecture requires reliable measurement. To assess reliability, we estimated twice per observer and found a high correlation between sessions (*r* = 0.96; **Figure 4A**). The data lie slightly above the identity line due to a small improvement across sessions (average of 10%). Similarly, we measured *A* twice by having two researchers independently draw the retinotopic map boundaries (from the same fMRI data) and found high agreement in surface area between the two researchers (*r* = 0.94, 0.88, 0.74, 0.73 for V1 to V4, respectively; **Figure 4B**). Although we did not have two separate scan sessions per observer, retinotopic mapping parameters in the early visual cortex are stable ac oss scans^37,38^, and the uncertainty in *A* from scan-to-scan variability is negligible compared to differences in how researchers use heuristics to draw map boundaries^39^.

**Figure 4.**
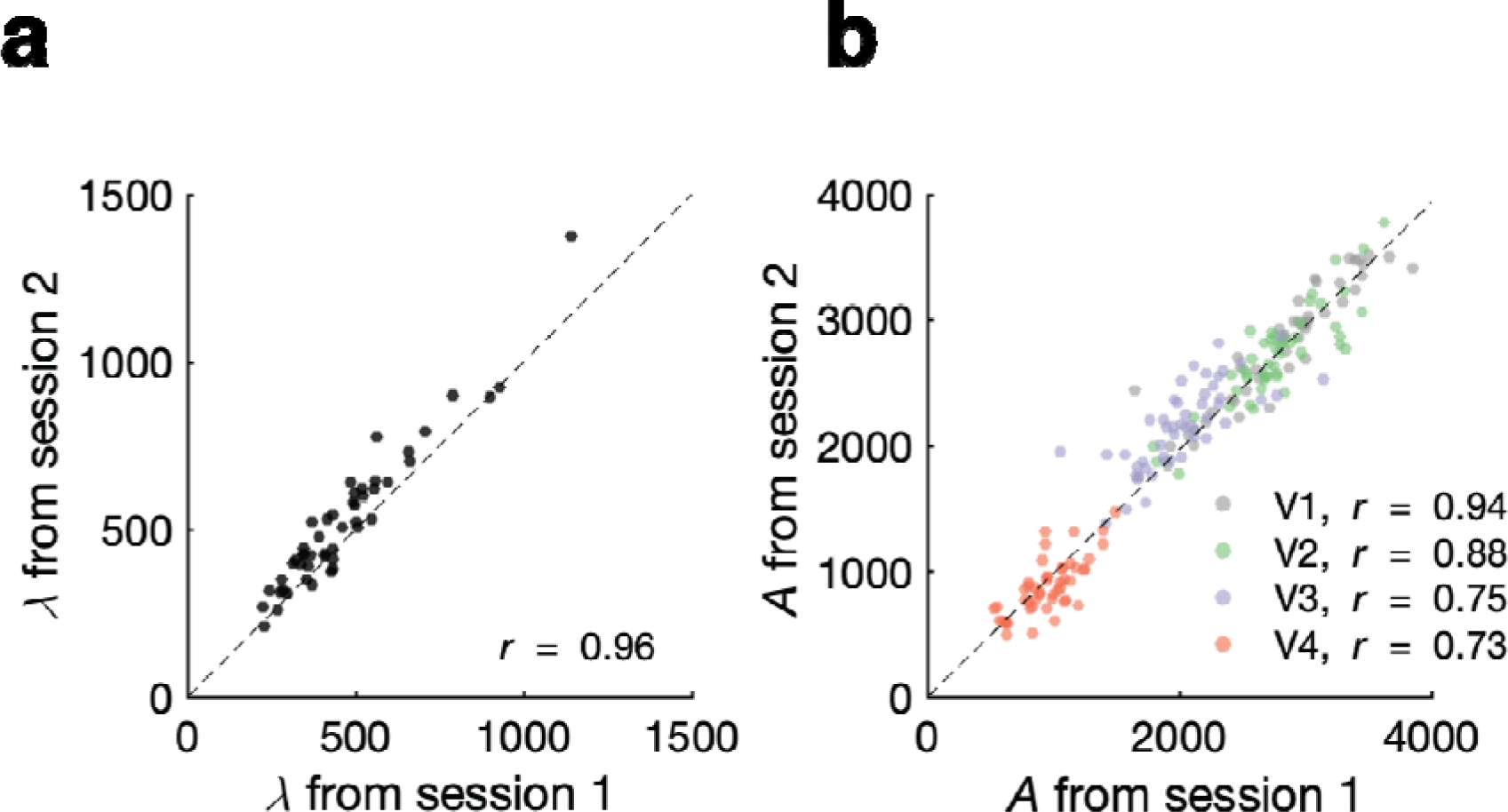
Measurement reliability of crowding distance and map size. (a) Test-retest correlation of. Each dot is one observer, with one value of measured from each of two test sessions on different days. (b) Correlation in map surface area across two researchers. Two researchers independently delineated the boundaries of the four retinotopic maps in each hemisphere of each observer. The surface areas plotted are bilateral (summed across L and R), and are limited to 10 deg of eccentricity.

### Cortical crowding distance is conserved in V4 but not V1, V2, or V3

Relating surface area to crowding, we plot each observer’s *A* vs. their, separately for V1 to V4 (**Figure 5a**). Only V4 shows high correlation between □ and *A* (*r* = 0.65; **Figure 5b**). In V1 to V3, there is no relationship between □ and *A*, as indicated by the practically zero correlation coefficients and the circular covariance ellipses in the scatterplots. The correlation of with the size of V4 but not V1 to V3 suggests that V4 size is independent of the sizes of the V1 to V3 maps. This is indeed the case in our dataset, with correlations between V4 size and V1, V2 and V3 of *r* ≤ 0.11 (**Supplementary figure 6**). Previous reports also found no correlation between the size of V1 and V4^39^, and little correlation between the size of V1 and V3^36^.

**Figure 5.**
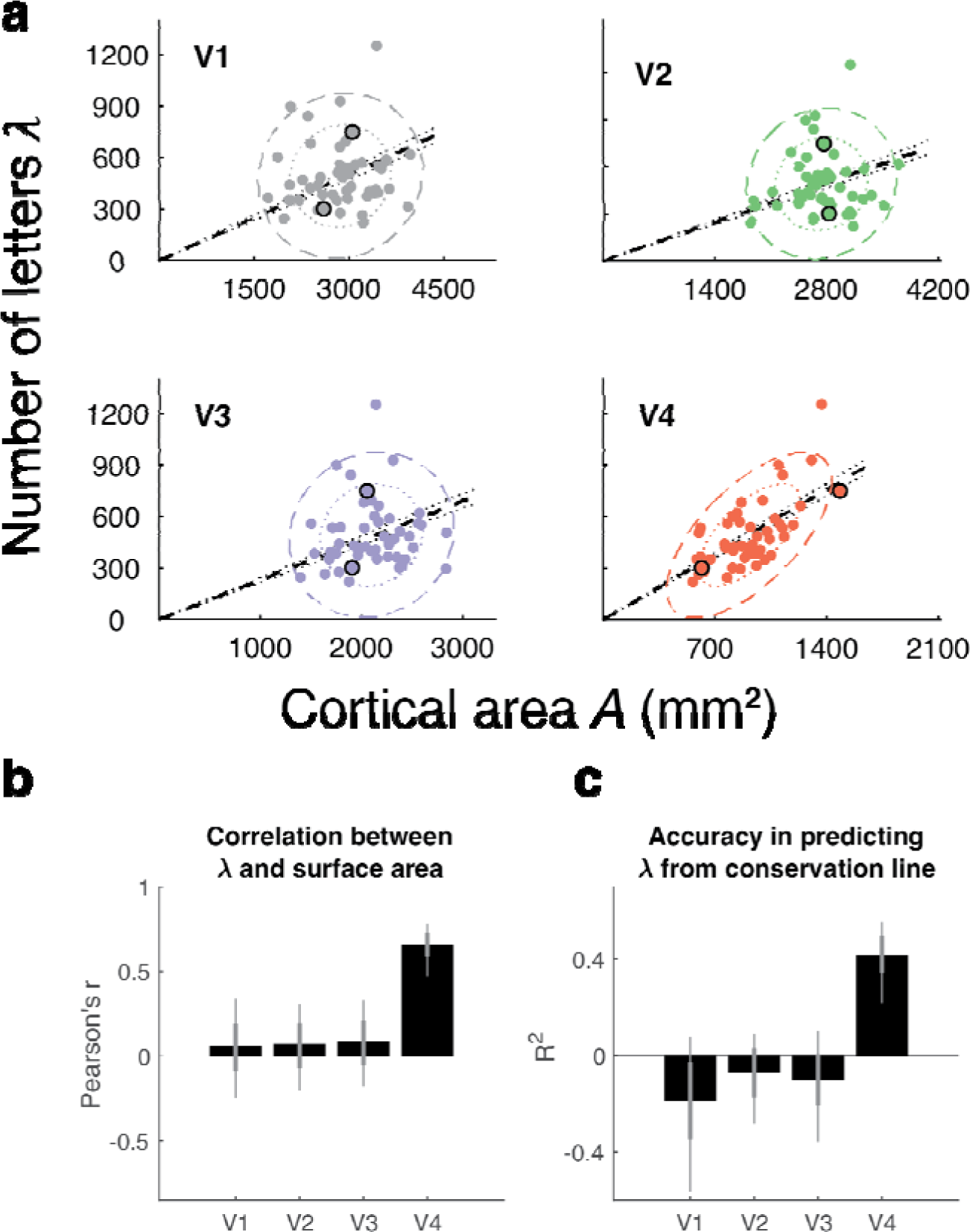
A test of cortical crowding distance conservation in V1 to V4 in 49 participants. (a) Each panel plots one point per observer as vs *A*. Both and *A* are computed for the bilateral visual field up to 10 deg eccentricity. Black outlines around two dots in each panel show data for the two observers from Figure 2. The colored ellipses are the 1-SD and 2-SD contours from the covariance ellipses. The black dashed lines are the fits by regression with one parameter, slope. Hence these lines are the predictions assuming cortical conservation of crowding. The dotted lines show 68th percentile of the conservation fit. (b) The correlation coefficient *r* between and *A* for V1 to V4. (c) The coefficient of determination R^2^ by assuming conservation (dashed lines in panel a). For panels b and c, we bootstrapped the data 10,000 times across observers; the bars are the medians and the error bars the 68% (thick) and 95% (thin) confidence intervals across bootstraps. See supplementary Figure 2 for results split by which of the two researchers defined the cortical maps, and supplementary Figure 3 for results split by hemisphere.

If a map conserves cortical crowding distance across observers, not only will □ and *A* be correlated, but the two measures will be related by a scale factor with no offset (**Eq. 1**). The V4 data are close to scaling, indicated by the orientation of the covariance ellipse being close to the conservation line in **Figure 5a**. We quantified how well the conservation hypothesis explains variation in by fitting the scale factor *k* in equation 1. We find that the conservation conjecture for V4 surface area accounts for 43% of the variance, whereas for V1 to V3 it accounts for none (**Figure 5c**).

The slope of the conservation line (dashed line in **Figure 5a**) allows us to estimate the required spacing between letters on the V4 map. We do so by re-expressing **Equation 1** in terms of the cortical crowding distance *c*, which measures center-to-center threshold spacing of letters in mm on the cortex:

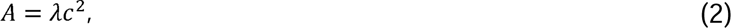

where

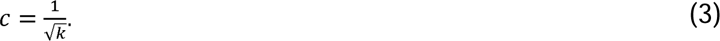

For V4, the slope *k* is 0.54 letters per mm^2^ [CI 0.52 - 0.57] which implies that the cortical crowding distance *c* is 1.36 mm [CI 1.33 - 1.39]. According to the conservation model, this single number, 1.4 mm, is the required letter-to-letter spacing, despite very different map sizes and crowding distances across observers. This estimate of *c*, 1.4 mm, is robust to variations in the assumed parameter values used to compute *λ* (**Supplementary Analyses 1 and 2**). Moreover, the value of *c* obtained here is close to the value that we obtained in a pilot experiment (1.25 vs. 1.36; **Supplementary Figure 5**). There are only three common participants between the pilot and the main experiment.

## Discussion

Our main finding is that despite large individual differences in psychophysical crowding distance and map surface area, crowding distance is conserved on the V4 map. We concluded this by comparing the number of letters that fit into the uncrowded visual field *λ* and the surface area of retinotopic maps *A*, finding that *λ* scales with *A* for V4 but not for V1 to V3. A model assuming scaling explains over 40% of the variance in *λ* for V4, and none for V1 to V3. The scaling implies a cortical crowding distance in V4 of 1.4 mm.

### Why is cortical crowding distance conserved?

We speculate that this 1.4-mm crowding distance is conserved across observers because there is a computational unit composed of a fixed number of V4 neurons needed to isolate an object. A fixed neural count implies a fixed area, which we estimate to be *A*/*λ* = 2 mm^2^ of cortex. This area contains about a quarter million neurons^40^. This proposed V4 computational unit isolating an object is analogous to a V1 hypercolumn analyzing local scale and orientation^41^, about 1.5 mm^2^ in human (and 0.6 mm^2^ in macaque)^42^.

The V1 hypercolumn is a well-defined concept. Functional domains have been identified in V4 (reviewed by Pasupathy et al^43^), but are not sufficiently well defined to measure their size. Nonetheless, an indirect measure of cortical organization, the spatial extent of direct lateral connectivity, has been quantified in macaque: Tracer studies estimate that 95% of intrinsic horizontal connections in V4 are within 1.9 mm, and 80% within 1.4 mm^44^, close to our estimated V4 cortical crowding distance.

A related interpretation is that the spacing limit for recognition relates to receptive field size rather than cortical distance. In this interpretation, if multiple letters are within the same receptive field, the representations of the separate objects would be degraded, as suggested in biased competition theory^45^. The two possibilities are not mutually exclusive. Receptive field size tends to be inversely correlated with map size, at least in human V1^31^ and so spacing and receptive field size are likely to make similar predictions. Here we focused on area rather than receptive field size because map size is a robust, reliable measurement, whereas receptive field size estimates have considerable uncertainty, both in human neuroimaging^46^ and single unit electrophysiology^47^.

### Measuring anatomy vs. function

Physiological methods have linked crowding to a variety of visual maps, with no clear consensus that a single visual area is closely linked to crowding distance (**Table 1**). Physiological studies of crowding, and computational models based on physiology, tend to compare neural responses to a target with different crowding conditions, such as small vs large spacing^12,22,23^. When the stimulus changes, neural responses also change. As a result, the particular metric calculated from the neural responses by the experimenter, such as target decodability from a set of sampled neurons^22^ or BOLD response magnitude, is likely to be affected in most visual areas. This creates ambiguity in identifying a single brain area as the crowding bottleneck. This is unsurprising since object recognition relies on computations spanning much of the visual system^48^.

In contrast, our search for a neural basis of crowding measured anatomy rather than physiology and exploited individual differences, measuring both crowding and map size in the same 49 individuals. An advantage of anatomical measures such as map size is that they are robust and do not depend on task, stimulus, and attentional state (for example, see Extended Figure 9 in ref [^49^]). Unlike the physiological approach, we find a sharp divide: no relationship between crowding psychophysics and map size in V1 to V3, and a strong relationship in V4.

### V4 as a gateway to the ventral stream

The sharp divide between the results in V4 and those in V1 to V3 surprised us. Why should the neural resources in a single brain area be linked to crowding thresholds, when image recognition involves many stages of processing, from optics to retina to the brain? We interpret this sharp divide as evidence for a spacing bottleneck in recognition. Suppose that each visual area has its own spacing threshold, such that neural responses to a target object are suppressed whenever there are similar objects closer to the target than its threshold separation. Our interpretation is that the *psychophysical* crowding distance is determined by the visual area in which the separation threshold is largest, meaning the area most sensitive to stimulus spacing. Our study implies that among posterior maps V4 is the most sensitive, i.e., has the largest spacing threshold. This result is consistent with known properties of visual maps. For example, from V1 to V4, the size of receptive fields increases^50–52^ and the surface area decreases^29,39,53^, so that there are decreasing numbers of neurons to analyze a given part of the visual field in higher visual areas. Hence if objects are sufficiently spaced for V4 (greater than about 1.4 mm), they will also be sufficiently spaced for earlier areas such as V1 to V3. The opposite, however, is not true: if stimuli are sufficiently spaced for V1, they can still be insufficiently spaced for V4. This is why visual areas prior to V4 are not spacing bottlenecks and do not correlate with crowding measures.

This argument still leaves open the question of why V4 size should be a spacing bottleneck rather than an even later stage of cortical processing. Indeed, there is spatial sensitivity in ventral temporal brain areas anterior to V4^39,54–56^. Nonetheless, V4 is likely one of the final processing stages prior to visual information being analyzed by specialized regions such as those involved in recognizing words^57^, faces^58^, and places^59^. Hence if object spacing is sufficient for processing in V4, then recognition by downstream areas is possible. This conclusion is consistent with connectivity analyses suggesting that V4 acts as a gateway to the ventral stream recognition areas^43,60^. It is possible that the size of other, downstream ventral temporal regions would also correlate with crowding behavior, perhaps as strongly as, or more strongly than, V4. The relationship between crowding and cortical size is more difficult to study in these areas, however, as retinotopic mapping is less precise in anterior maps, and the criteria for defining boundaries, and even the number of discrete areas, remains uncertain for category-selective regions ^61^. Thus, we can conclude that V4 is the *first* visual area to show a strong relationship to crowding behavior, but we cannot draw conclusions about whether it is the only such area.

Our proposal that V4 acts as a spacing bottleneck for recognition is consistent with several properties of crowding. First, crowding distance is similar for many types of stimuli (letters, faces, other objects)^62^, which seems inconsistent with crowding arising independently in areas that are selective to different kinds of object. Second, the nonlinearity in the growth of crowding distance with eccentricity^63^ matches the nonlinear decline of cortical magnification with eccentricity in human V4 map^64^. This nonlinearity is not present in V1, V2 and V3^36^.

### Homology between human and macaque V4

Our results reveal a tight link between the size of the human V4 map and psychophysical crowding distance. In arguing that V4 mediates crowding, we referred to literature on both macaque and human V4. However, the extent of homology between human and macaque V4 remains uncertain^65^. In both species, V4 has been identified as part of the ventral visual processing stream, with an important role in recognition^53,66^. On the other hand, human V4 is located on the ventral surface of the occipital lobe^53,67,68^ (but see Refs ^69,70^), whereas macaque V4 is split into a ventral and dorsal part^52^. Nonetheless, macaque recognition, like human recognition, is limited by crowding^23,71^, and it is plausible that V4 acts as a bottleneck in both species. Whether or not the surface area of macaque V4 predicts individual differences in macaque crowding thresholds requires independent validation.

### The signal and the noise

The reader may share our curiosity about the relationship between *λ* and A at a finer-grained scale than we reported. Unfortunately, splitting the data into small parcels decreases the signal-to-noise ratio. For example, the test-retest correlation of *λ*, pooled across the four meridians, was 0.96, whereas the test-retest correlations for the left and right meridians alone were 0.84 and 0.71, respectively (Supplementary Figure 4). The lower reliability of the single meridian behavioral estimates reduces the power to detect a relationship with cortical measures. Nonetheless, the main pattern of results holds for various subsets of our data (Supplementary figures 2 and 3), albeit with weaker relationships. For example, the correlation coefficients between *λ* and *A* for only one V4 hemisphere and one meridian are *r*=0.58 and *r*=0.30 (right hemisphere/left meridian, and left hemisphere/right meridian), compared to *r*=0.65 bilaterally.

What limits the precision our measurements? Retinotopy with fMRI is effective but imperfect^38,72,73^. There are many non-neural factors contributing to the fMRI signal^74^. For example, vascular artifacts could cause a portion of a map to have poor signal and be excluded from our estimate of *A*, particularly in the left hemisphere of V4^75,76^. Also, some participants have unusual V2 or V3 maps^77^, which might not be well captured by our manual delineation procedures. Such artifacts can introduce systematic bias at the level of the individual subject, meaning both researchers underestimate or both overestimate *A*. However, as fMRI artifacts are unlikely to be related to behavior, individual biases translate to random (unbiased) noise at the level of correlation across the group. Random noise puts an upper limit on goodness of fit but has little effect on estimate of slope.

## Conclusion

We exploited individual differences to uncover a strong connection between visual crowding and V4 size. Individual differences can help characterize visual mechanisms^78^, and they are important in their own right. Extreme individual variation becomes a disorder, as in dyslexia, dyscalculia and apperceptive agnosia, which are associated with unusually large crowding distances^5–7^. Our discovery that the size of the V4 map predicts crowding might help understand these disorders and is a step in settling the long-standing dispute about the cortical locus of crowding.

## Methods

### Participants

Fifty participants (32 females, 18 males, mean age 27 years) were recruited from New York University. All participants had normal or corrected-to-normal vision. The participants completed one fMRI session to measure retinotopic maps (45 min) and two behavioral sessions to measure crowding distance (30 min each). One participant was dropped from the analysis due to incomplete fMRI data which did not allow for reliable estimation of retinotopic maps. All participants provided written informed consent and approved the public release of anonymized data. The experiment was conducted under the Declaration of Helsinki and was approved by the NYU ethics committee on activities involving human participants. Data from one observer was excluded because the fMRI session was not completed.

### Pilot study

We conducted a pilot study with 26 participants and reported the results in a conference abstract^79^. The new experiments reported here were conducted to include test-retest measures, obtain a larger number of participants, and assess replicability. Eight of the pilot subjects were re-tested in the main experiment including both new crowding data and new retinotopic mapping. We performed the same analysis of the pilot data as presented in the main text. The results reported in the pilot study are largely independent and highly consistent with those reported in the main text (**Supplementary Figure 5**).

### fMRI stimulus display

Participants viewed a retinotopic mapping stimulus in the MRI scanner using a ProPixx DLP LED projector (VPixx Technologies Inc., Saint-Bruno-de-Montarville, QC, Canada). The stimulus image was projected onto an acrylic back-projection screen (60 cm × 36.2 cm) in the scanner bore. The projected image had a resolution of 1920L×L1080 and a refresh rate of 60 Hz. The display luminance was 500 cd/m^2^. Participants viewed the screen at 83.5 cm (from eyes to the screen) using an angled mirror that was mounted on the head coil.

### fMRI Stimuli

The mapping stimulus was generated in MATLAB 2017a and was presented using the Psychophysics Toolbox v3^80^ and custom vistadisp software (https://github.com/WinawerLab/vistadisp) on an iMac computer. For 27 participants, stimulus images were shown within a sweeping bar aperture, identical to the stimulus in Himmelberg et al^37^. For 33 participants, the stimulus image patterns were shown within two types of apertures in alternative scans, a sweeping bar and a wedge+ring. These stimulus apertures were similar to the ones used in the NSD dataset^49^. Unlike NSD, our stimuli changed location in discrete steps and had a larger maximum eccentricity. In short, there were six runs of retinotopic mapping, each lasting 5 minutes (300 TRs of 1 sec). In recent work we found that the inclusion of the wedge+ring aperture was important for getting reliable estimates of pRFs centered near the fovea^37^. Each aperture type was shown three times in an interleaved order. The carrier image pattern was presented with a 3 Hz temporal frequency and was composed of colorful objects, faces, and scenes at multiple scales. The faces, objects and scenes were made for a study of object recognition^81^, and then later incorporated into retinotopic mapping, with the addition of an achromatic pink-noise (1/f) background^82^. The motivation for including faces, objects and pink noise was that a more varied carrier pattern might effectively drive responses across visual cortex, especially higher-level areas The stimulus image pattern was windowed within a circular aperture (12.4° of maximum radius). The apertures were superimposed on a polar fixation grid placed upon a uniform gray background, with a red or green dot at the center (3 pixels, or 0.07°). Participants completed a fixation task to ensure that they were maintaining central fixation and remained alert throughout the scan. Participants were required to respond, via button press, when the central fixation dot changed from green to red, or vice versa.

### Data acquisition and preprocessing

Structural and functional data were acquired on a 3T Siemens MAGNETOM Prisma MRI scanner (Siemens Medical Solutions, Erlangen, Germany) at the Center for Brain Imaging at NYU. Both the structural and functional images were acquired using a Siemens 64-channel head coil. MPRAGE anatomical images were acquired for each participant (TR, 2400 ms; TE, 2.4 ms; voxel size, 0.8mm^3^ isotropic; flip angle, 8°) and were auto-aligned to a template to ensure a similar slice prescription for all participants. Six functional echo-planar images (EPIs) were collected from each participant using a T2*-weighted multiband EPI sequence. The parameters were as follows: repetition time (TR) of 1000 ms, echo time (TE) of 37 ms, voxel size of 2 mm³, flip angle of 68°, multiband acceleration factor of 6, and posterior-to-anterior phase encoding. Additionally, two distortion maps were acquired to correct susceptibility distortions in the functional images: one spin-echo image with anterior-posterior (AP) and one with posterior-anterior (PA) phase encoding. Anatomical and functional preprocessing was performed using fMRIPrep v.23.1.2^83^. For each participant and each run, fMRIprep produced a BOLD time series on the participant’s native FreeSurfer surface.

### Population receptive field model

We first averaged the time-series data from the three repetitions of bar aperture and three repetitions of wedge and ring apertures resulting in two functional time series. For the 27 participants who saw images only through a bar aperture, we averaged all 6 runs. We used these average data to fit the pRF model. The analysis was conducted using Vistasoft (https://github.com/WinawerLab/prfVista;). A pRF is modeled as a circular 2D-Gaussian, as described in Dumoulin and Wandell^84^. The Gaussian is parameterized by values at each vertex for x, y, and σ. The x and y parameters specify the center position of the 2D-Gaussian in the visual field. The σ parameter, the standard deviation of the 2D-Gaussian, specifies the size of the receptive field. The 2D-Gaussian is multiplied pointwise by the stimulus contrast aperture (apertures were prepared at a resolution of 101 pixels × 101 pixels), and then convolved with a hemodynamic response function (HRF) to predict the BOLD percent signal change. The HRF is parameterized by 5 values, describing a difference of two gamma functions, as used previously^84^. The HRF was assumed to be the same across vertices within a participant but differed among participants. We use a two-stage coarse-to-fine approach described in detail by Dumoulin and Wandell^84^, and the addition of the HRF fit is described in detail by Harvey and Dumoulin^31^

### Defining the size of visual areas (V1-V4)

Two authors, JWK and BSQ, independently defined ROIs (V1-V4) by hand using neuropythy (https://github.com/noahbenson/neuropythy)^85^. To define visual maps, researchers followed the heuristics of Wandell^86^ for V1-V3, and of Winawer and Witthoft for V4^87^. Note that while we have developed automated tools for defining visual maps, our tools have only been validated for V1 to V3^73^. To match the maximum eccentricity of the psychophysical stimuli we restricted all maps to a maximum eccentricity of 10 deg. The size of each map was calculated as the sum of the surface areas of all vertices included in the map on the mid-gray cortical meshes, defined by FreeSurfer as the surfaces hallway between the meshes at the gray/white interface and the pial/gray interface. Finally, each map surface area was summed across hemispheres.

### Measuring crowding distance

Data were acquired with the CriticalSpacing software^88^, allowing for reliable and relatively fast measurement of crowding distances. Our crowding database consists of measurements of crowding distance with the Sloan font with radial flankers. We measured crowding at 8 different peripheral locations in the visual field: 2 eccentricities (5, 10 deg) along the four cardinal meridians (upper, lower, left and right). Each observer participated in two crowding sessions (each lasting about 40 min). Sessions were separated at least by a day with a maximum of five days. In session one first block showed stimuli at 5 deg and the second showed stimuli at 10 deg. In the second session the order was reversed.

### Crowding stimulus display

Each testing session was completed on an Apple iMac 27” with an external monitor for stimulus presentation. Screen resolution was 5120 × 2880. Apple iMac has AMD graphics for optimal compatibility with the Psychtoolbox imaging software. The Systems Preference: Displays: Brightness slider was set (by calling MacDisplaySettings.m in the Psychtoolbox) to 0.86 (range 0 to 1) to achieve a white background luminance of about 250 cd/m^2^. The observer viewed the screen binocularly at 40 cm distance from the screen. Stimuli were rendered in MATLAB 2021 using the Psychtoolbox^80^

### Crowding stimuli

To measure crowding we show a three-letter trigram. For each trial, the three letters are drawn randomly, without replacement, from the 9 possible letters. Letters are rendered as black in the Sloan font, presented on a uniform white background of about 250 cd/m2. For the Sloan font we use the letters “DHKNORSVZ” as possible targets and flankers. For crowding, each trigram is always arranged radially. Each testing session is about 30 minutes, measuring crowding for targets at two eccentricities, 5° and 10°. The eccentricities are measured in separate blocks of about 15 min each. Each block measures crowding thresholds in the four cardinal directions, with four interleaved staircases. Fixation is displayed until the observer presses a key to initiate the trial. Then the letter trigram appears on the screen for 150 ms and the computer waits for the observer to identify the middle letter by choosing the letter using a mouse click. Letter choices appear on the screen after stimulus offset. Observers are instructed to return their eyes to fixation before clicking their response. The computer waits indefinitely for each response. 300 ms after the observer’s response, the next trial begins as soon as the observer fixates the center of the crosshair for 250 ms.

### Eye tracking

For all thresholds, we controlled for eye movements. To measure fixation, we used an Eyelink 1000 eye tracker (SR Research, Ottawa, Ontario, Canada) with a sampling rate of 1000 Hz. To allow for short viewing distance (40 cm) we used their Tower mount setup with a 25 mm lens mounted on the Eyel nk camera. We allowed participants to move their eyes within a 1.5-degree radius from fixation. We checked eye position during the stimulus presentation. If during the stimulus presentation, observers moved their eyes more than 1.5 deg away from the fixation, the fixation cross turned red, and observers were asked to repeat the trial. Each threshold was estimated based on 35 trials with correct fixation. On average, 10% of trials had to be repeated.

### Measuring crowding distance

Pelli’s^88^ procedure estimates the crowding distance. Letter spacing is controlled by QUEST. Spacing scales with letter size, maintaining a fixed ratio of 1.4:1. We set the guessing rate parameter gamma to the reciprocal of the characters in the test alphabet for that font, usually 9. We set the “finger error” probability delta to 1%, so that QUEST can better cope with an occasional reporting mistake. At the end of each block, QUEST estimates the threshold (crowding distance). We devote 35 trials to measure the threshold for a condition. In each block we randomly interleaved the four meridian locations, to get a threshold for each condition, while keeping the observer uncertain as to which condition is present on each trial.

### Calculating the Bouma factor *b*

For each participant we measured 16 crowding distances (2 eccentricities × 4 cardinal meridians × 2 sessions). For each session, crowding distances were converted to Bouma factors by dividing them by the target eccentricity. (This calculation omits because it’s negligible, 0.24 deg << 2.5 deg.) Next, we estimated one Bouma factor for each session by taking the geometric mean across all 8 visual field locations. The final estimate for each observer was calculated as the geometric mean of Bouma factors from sessions 1 and 2.

### Calculating the number of uncrowded letters

We used the Bouma law to calculate the number of letters that fit into the visual field without crowding. The Bouma law states that the radial crowding distance *s*_r_, in deg, grows linearly with eccentricity, in deg,

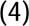

where and are fitted constants. Tangential crowding distance *s*_t_ is generally found to be proportional to radial crowding distance,

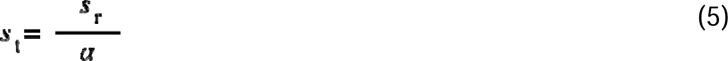

where (typically 2) is the ratio of radial-to-tangential crowding distances^26^.

When letters are packed as tightly as possible without crowding, the local density, in letters per deg2, is the reciprocal of the product of the radial and tangential crowding distances. The number of uncrowded letters in the visual field is the integral of that density over the visual field,

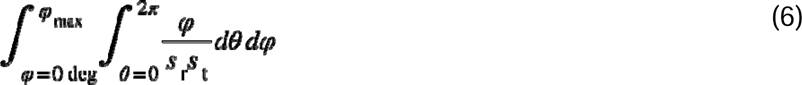

where is the eccentricity (in deg), is the maximum extent of the visual field (in deg), is the polar angle, and and are the radial and tangential crowding distances (in deg). Integrating yields a formula,

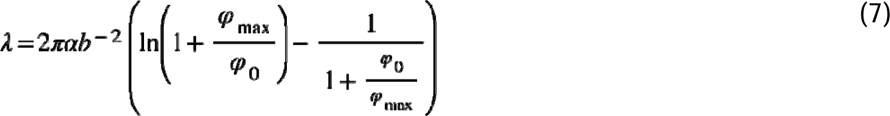

The last term is an approximately logarithmic function of the ratio _max_/ _0_ (**Supplementary Analysis 2**). We set _max_ = 10 deg, corresponding to the maximum eccentricity of both the psychophysical and retinotopic data. We took measured values, _0_ = 0.24 deg and = 2, from our prior study^63^. With these values, the formula reduces to

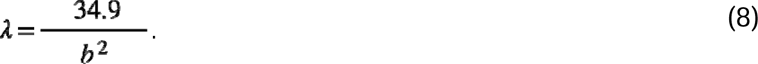

Our measure of interest, *c*, the cortical letter spacing at threshold separation, depends on. We analyzed the sensitivity of to the values of and (**Supplementary Analysis 1**).

### Relationship between *A and*

We quantified the relationship between *A*, cortical map surface area, and, number of letters in the uncrowded visual field, in two ways. First, we computed the correlation (Pearson’s *r*) and covarianc between the two variables. These calculations treat the two variables symmetrically. We visualize the covariance ellipses at 1-SD and 2-SD, scaling the x and y axes so that one standard deviation of each variable is the same length (**Figure 3a**). This ensures that the covariance ellipses are circular when the correlation is 0. Second, to explicitly test for conservation, we fit a linear model to *A* (independent variable) and (dependent variable), with the intercept set to 0, and asked how much of the variance in is explained (**Figure 3c**). Note that the variance explained can be negative, because a line through the origin may be a worse predictor (larger residuals in) than just assuming the mean (a horizontal line). For both the correlation coefficient and the variance explained, 68% confidence intervals were calculated by bootstrapping participants with replacement (*n* = 10000).

### Bootstrapping and confidence intervals

We quantified variability in the data with confidence intervals derived from bootstrapping over participants. Bootstrapping has the advantage over parametric statistical measures in that it does not make assumptions about the shape of the distributions. In a normal distribution, 68% of the data fall within one standard deviation of the mean, and 95% within two standard deviations. Hence by analogy, we report 68% and 95% CIs from bootstrapping. While we do not report formal null hypothesis significance tests (their scientific value is controversial^89,90^), the provided CIs allow for such inferences: when the 68% CIs of two groups do not overlap, this is similar to rejecting a null hypothesis that the two groups do not differ with an alpha parameter of 0.05. When the 95% CIs do not include 0, this is similar to rejecting the null hypothesis that the measure does not differ from 0 with an alpha of 0.05.

## Code and data availability

Code to produce figures and raw crowding data together with MRI surface files are deposited on GitHub. fMRI dataset in BIDS format is available on OpenNeuro (https://openneuro.org/datasets/ds005639/versions/1.0.0).

## Acknowledgments

Funding support from NIH NEI Grant R01-EY033628, NIH NEI Grant R01-EY027401, NIH NEI Grant R01-EY027964, NIH NEI Grant R01-EY031446, and NIH NEI core grant for vision science P30EY013079. The authors thank Professor Michael Landy for comments and discussion.

## Supplementary materials

**Supplementary Figure 1.**
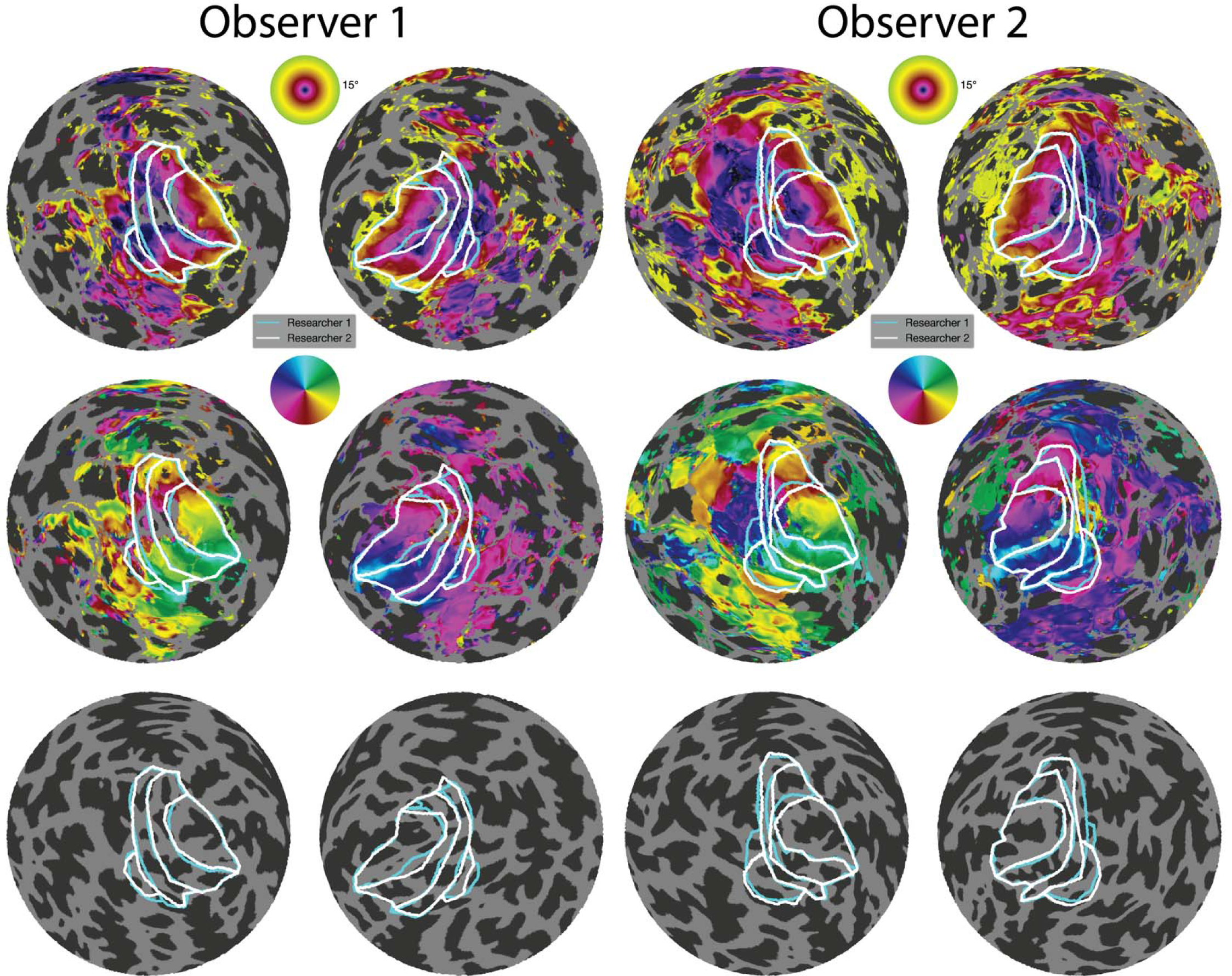
Retinotopic maps for Observers 1 and 2. For each observer we show flattened representations of their left and right hemispheres. The overlays show eccentricity (top), polar angle (middle) and underlying thresholded cortical curvature (bottom; dark regions are sulci, light regions are gyri). The two sets of outlines (blue and white) are the map boundaries drawn independently by two researchers. In rows 1 and 2, the color overlays are thresholded by pRF variance explained > 10%. Maps for all participants are available as images in Supplementary Data 1 (retinotopicMaps.zip).

**Supplementary Figure 2.**
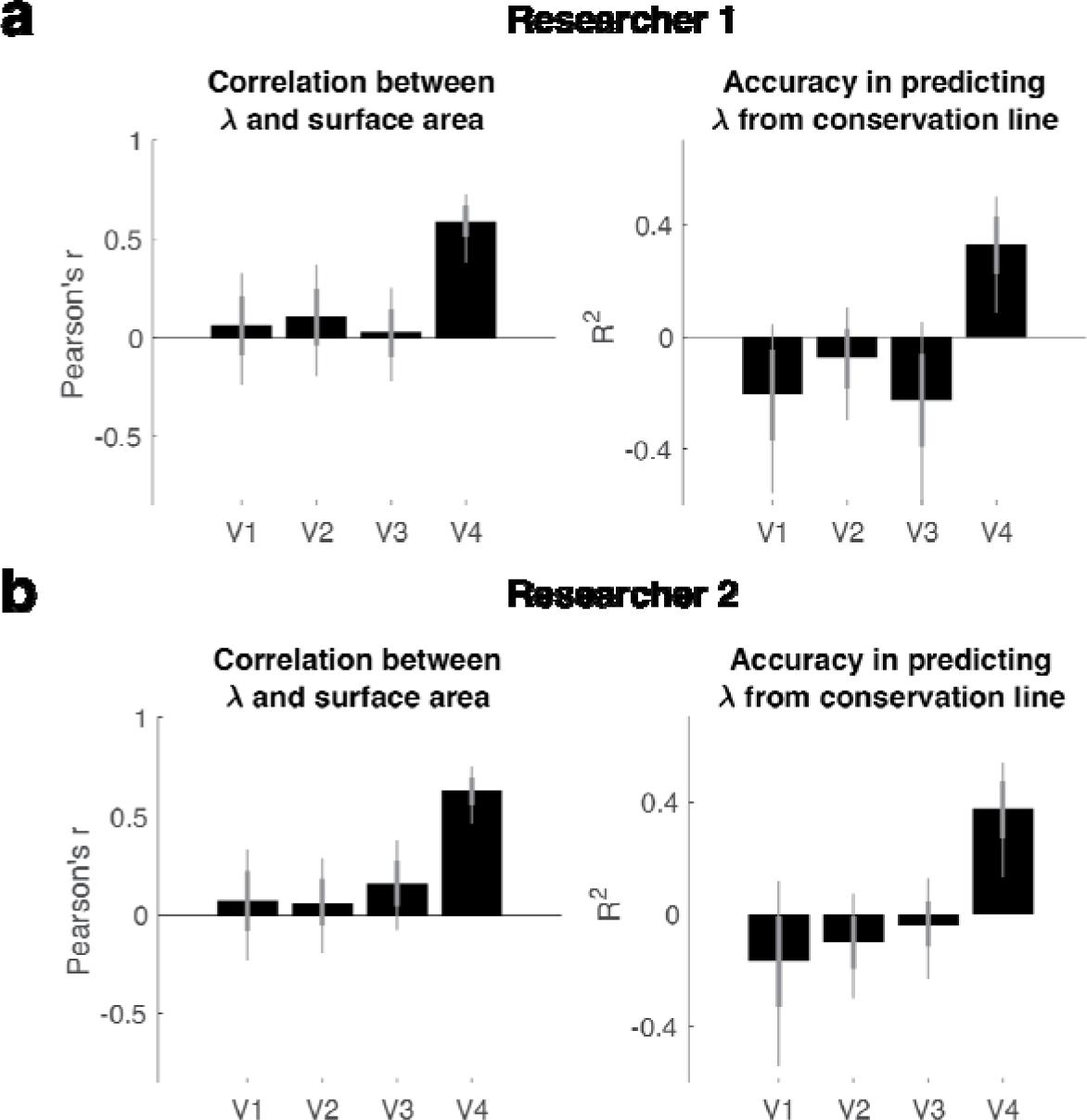
Model fits of vs *A* for V1 to V4, based on map boundaries from single researchers. Panel a shows the correlation coefficient (left) and variance explained from linear regression with 0 intercept (right) when using co tical map definitions from Researcher 1. Panel b is the same but for Researcher 2. The thick and thin error bars are 68% and 95% confidence intervals. The results can be compared to Figure 5b and 5c in the main text, which compute the same statistics for map surface area averaged across the two researchers.

**Supplementary Figure 3.**
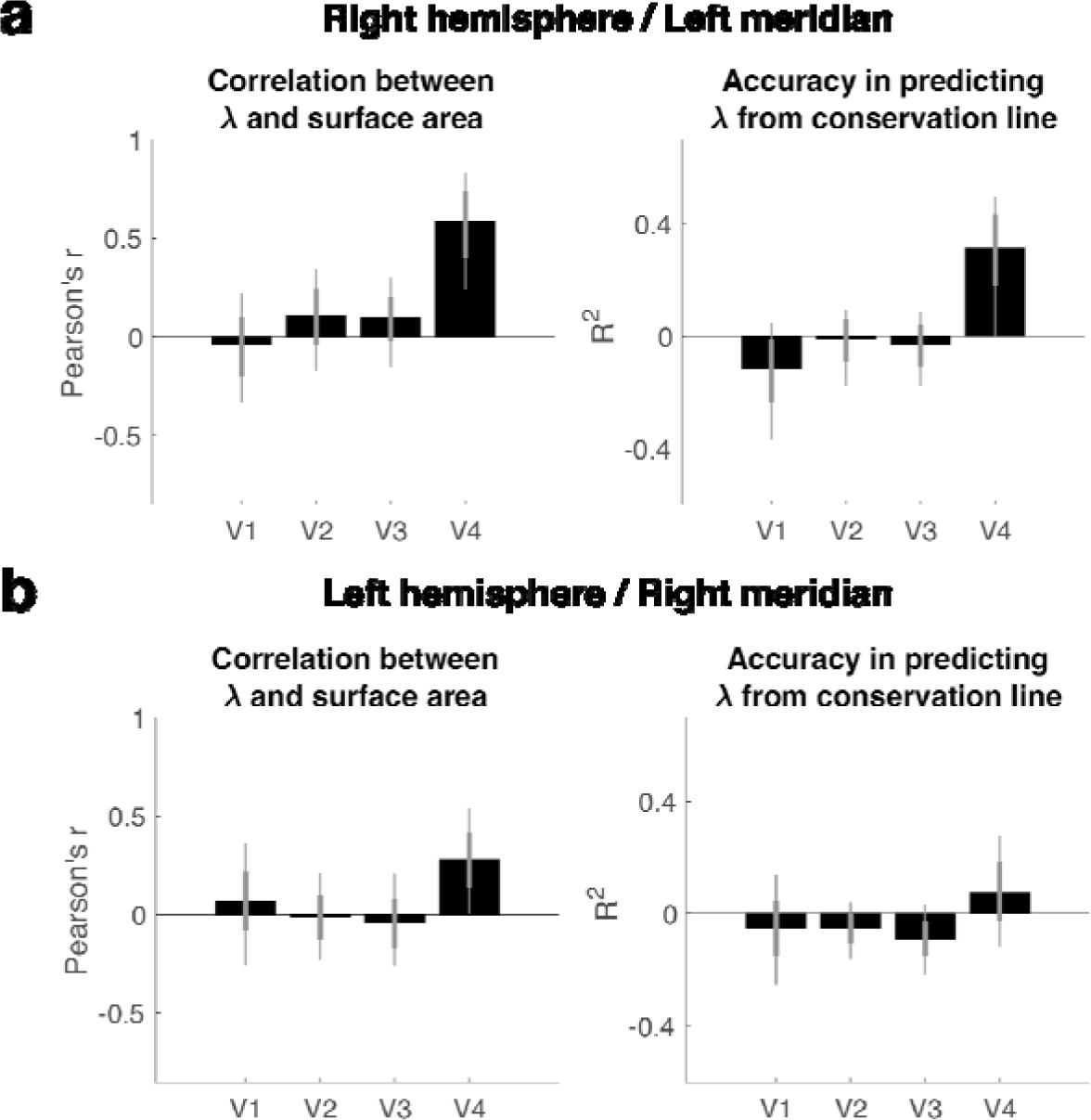
Model fits of vs *A* for V1 to V4, separated by cortical hemisphere. Both panels are as in Supplementary Figure 2, except that: (1) the map surface area is computed only for the right hemisphere (panel a) or left hemisphere (panel b) and is derived from crowding data only on the left horizontal meridian (Panel a) or right horizontal meridian (panel b). Surface area measures are averaged across the two researchers.

**Supplementary Figure 4.**
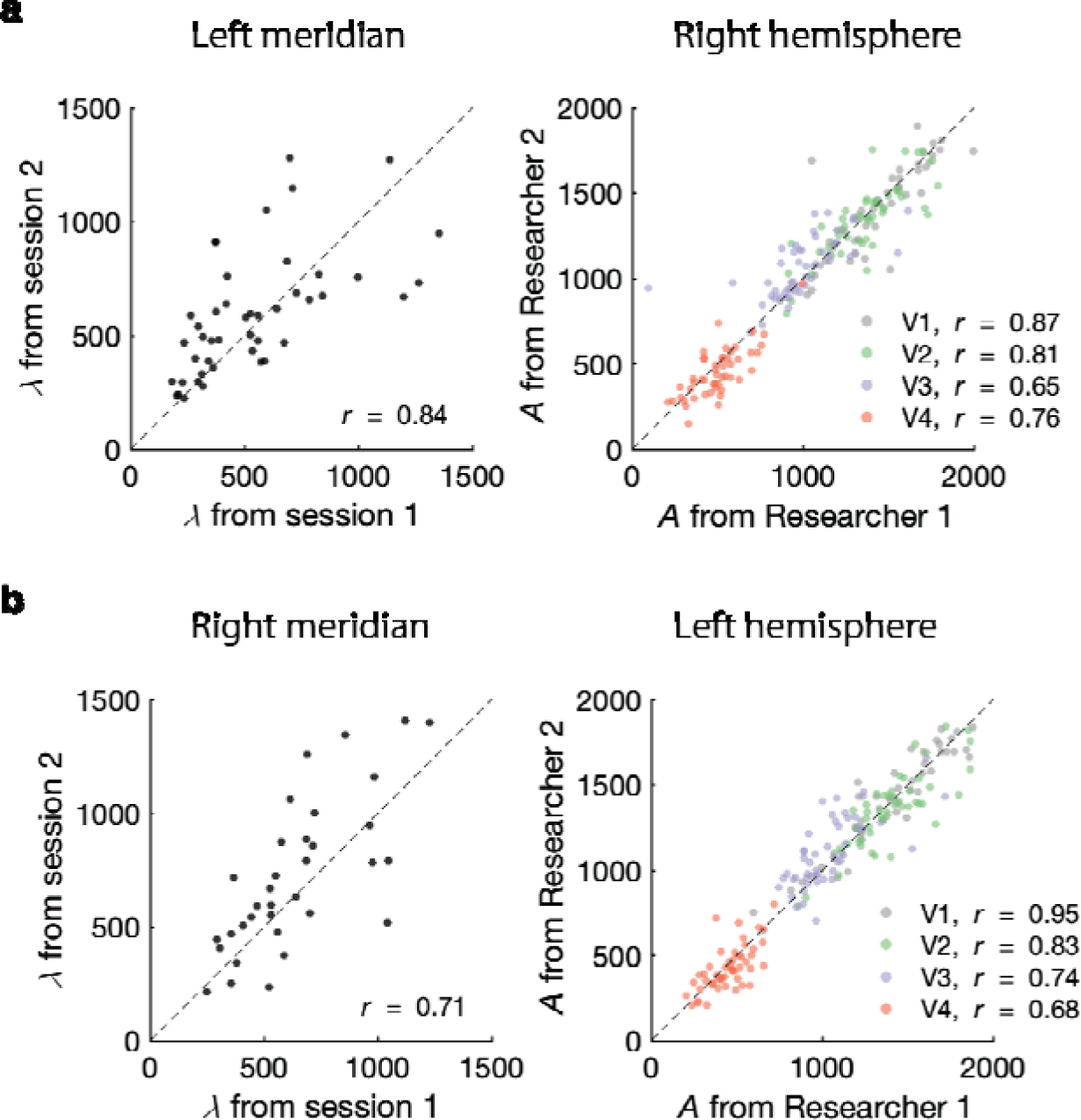
Measurement reliability of crowding distance and map size separated by hemisphere and visual field location. Both panels are the same as in Figure 4, except that panel a plots from the left meridian and right hemisphere, while panel b plots the data from right meridian and left hemisphere.

**Supplementary Figure 5.**
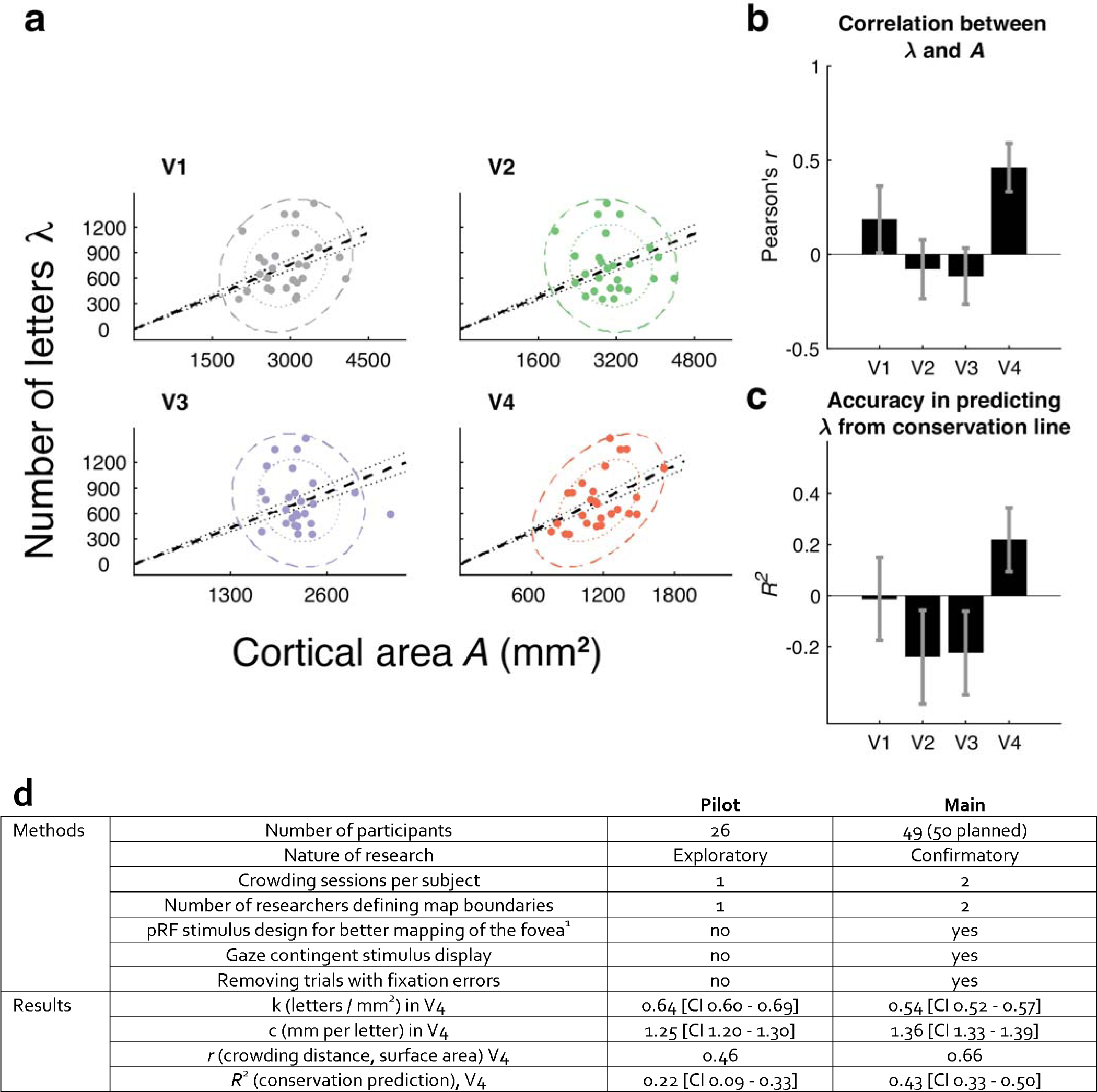
A comparison of pilot and main experiments. a, Number of letters *λ* vs cortical surface area A for the 26 observers in the pilot dataset. Otherwise plotted as in Figure 3a. b, The correlation between *λ* and *A* for V1 to V4 in the pilot dataset. Otherwise plotted as in Figure 3bc, Variance explained by conservation prediction. Plotted as in Figure 3c. d, A comparison of methods and results between the pilot and main experiments.

**Supplementary Figure 6.**
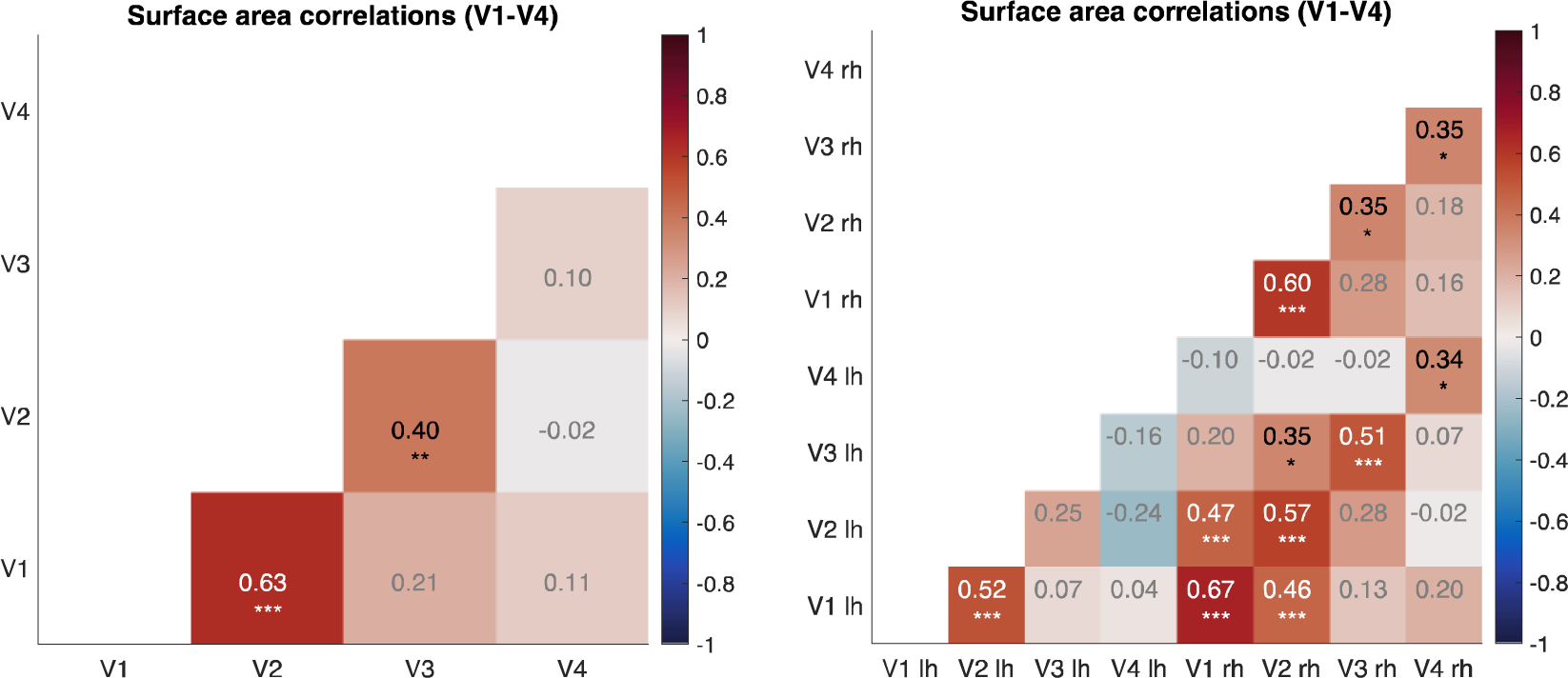
Correlations between surface areas of different maps. The left panel shows correlations for bilateral maps. Note that V4 has little correlation with the surface area of V1 to V3. The panel on the right shows the same but separated by hemisphere. The values in the cells are Pearson correlations (n=49). The asterisks indicate statistical significance of the null hypothesis test that the correlation between surface areas of two maps is exactly 0. One asterisk is p < 0.05, two asterisks is p < 0.01, and three asterisks mean p < 0.001.

**Supplementary Analysis 1: Sensitivity analysis of the estimated cortical crowding distance *c Effects of* α *and* φ_0_ on c.**

We concluded that cortical crowding distance is conserved on the V4 map, meaning that there is a single value for *c* (cortical crowding distance) across participants, estimated to be 1.4 mm. We estimated this value by regressing λ, number of uncrowded letters, on *A*, the surface area of the V4 map, assuming an intercept of 0 (eq 1 in main text), and then solving for *c* (equation 3 in main text).

Further, we reported uncertainty in *c* from bootstrapping across participants (Figure 3). Here we assess how c is affected by variation of λ across participants. The main-text calculation of λ assumed fixed values from our previous work^2^ for two parameters: the tangential-to-radial ratio a of crowding distances and the Bouma law intercept φ_0_. Here we calculate how the known parameter variations would affect the *c* estimate. We computed *c* 20,000 times, each time assuming different a values with φ_0_ fixed (10,000 calculations), or different φ_0_ values with a fixed (10,000 calculations). The values for the two parameters were based on the measured distributions, rather than assuming the mean as in the main text. From our prior study, the mean value of a was 2.10 (sd = 0.38), and the mean value of φ_0_ was 0.24 (sd 0.05). Supplementary Table 1 below reports the *c* median and 95% confidence intervals from the calculations.

**Supplementary Table 1.** Sensitivity of cortical crowding distance to variation in α and φ_0_.

|  |  |
| --- | --- |
| Median [95% CI] estimate of $c$, assuming:<br>$\alpha$ ($\square = 2.10$, $\square = 0.38$),<br>$\varphi_0 = 0.24$ | Median [95% CI] estimate of $c$, assuming:<br>$\alpha = 2.10$,<br>$\varphi_0$ ($\square = 0.24$, $\square = 0.05$) |
| 1.32 [1.14,1.64] | 1.36 [1.25,1.45] |

**Supplementary Analysis 2. *Effect of* _max_ on.**

The formula (Eq. 7) for number of uncrowded letters grows approximately logarithmically with _max_/ _0_ (**Supplementary Figure 7**, below). Our decision to measure (in psychophysics and fMRI) out to 10 deg eccentricity was somewhat arbitrary, but we can use equation 7 to consider how many more uncrowded letters might fit in the far periphery. As expected of logarithmic dependence, large changes in the maximum eccentricity produce only modest changes in the number of letters. For instance, doubling the maximum eccentricity _max_ from 10 to 20 deg would increase the number of uncrowded letters by only 13%, and increasing it tenfold, from 10 to 100 deg, would increase by 47%.

**Supplementary Figure 7.**
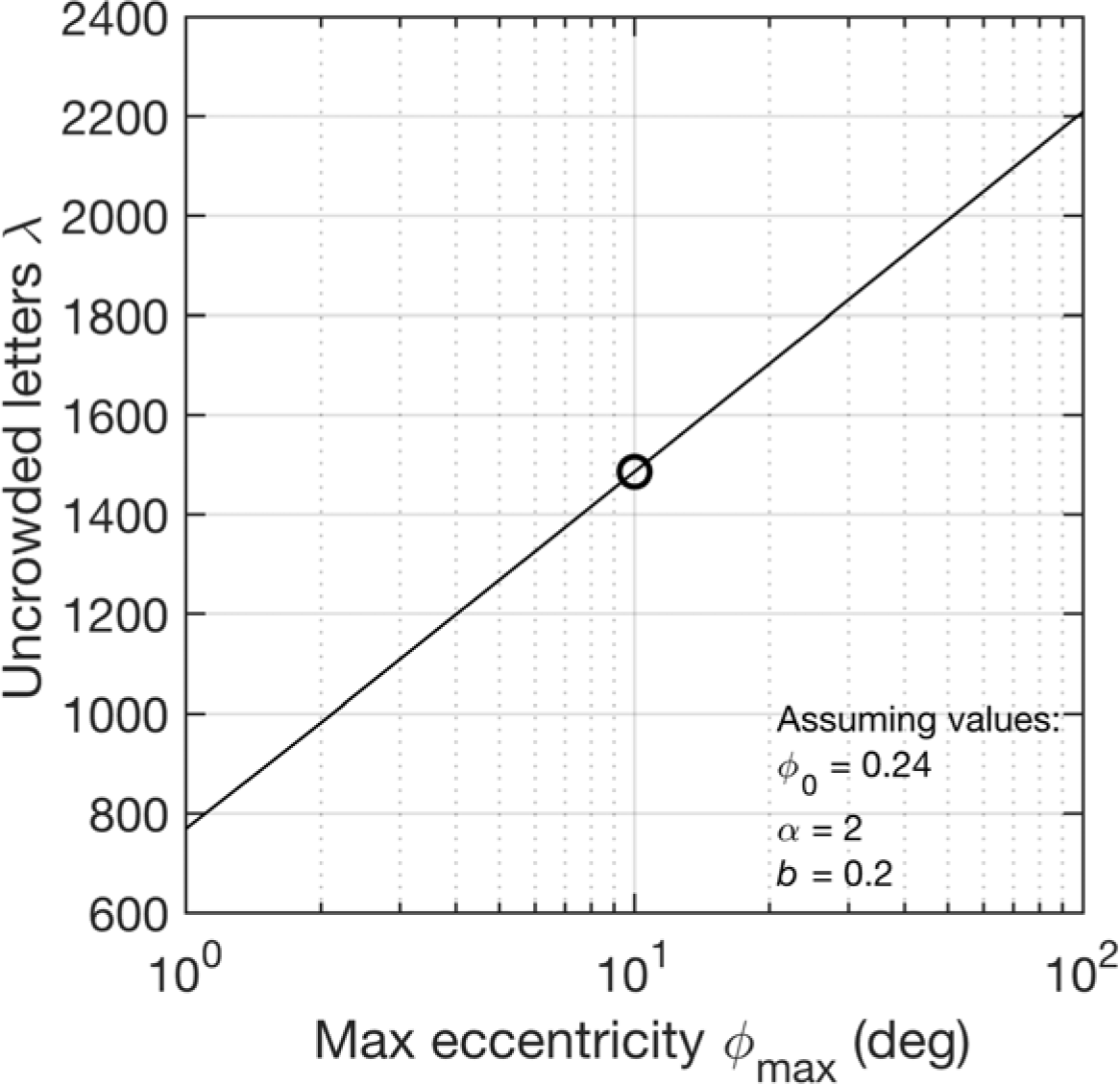
Sensitivity of number of uncrowded letters to maximum eccentricity max. Plots the formula (Eq. 7) for vs. max. This paper mostly sets max to 10 deg, shown by the circle. The nearly straight curve shows that the formula’s dependence on max is practically logarithmic.

**Supplementary Data 1**

Please see attached zip file which includes images of visual maps in all 49 subjects.

**Supplementary Data 2**

Please see attached csv file which includes a table of visual map sizes and crowding measures for all subjects.

